# The chromatin remodeler LET-418/Mi-2 regulates the intracellular pathogen response in the *C. elegans* intestine

**DOI:** 10.1101/2025.06.17.659900

**Authors:** Shweta Rajopadhye, Vladimir Lažetić, David Rodriguez Crespo, Emily Troemel, Peter Meister, Chantal Wicky

## Abstract

Chromatin remodeling provides essential transcriptional regulation for all biological processes. In *Caenorhabditis elegans,* the chromatin remodeler LET-418, a homolog of the human Mi-2β protein, plays a critical role in regulating development, organogenesis, tissue maintenance, stress resistance and lifespan. LET-418 is part of several chromatin remodeling complexes and contributes significantly to the balance between growth and defense mechanisms, yet its target genes remain unclear. Using DNA methylation profiling, we identified genomic binding sites and associated target genes of LET-418 and its MEC-complex-specific interactor MEP-1 in the intestine. Consistent with their presence in the same complex, the two proteins shared more than half of their target genes. Functional analysis revealed that LET-418 and MEP-1 target genes are highly active in the intestine and are involved in repressing innate immune responses, including the intracellular pathogen response (IPR). Consistently, in *let-418* mutants, IPR-induced genes, such as *pals-5* or *pals-2* are strongly upregulated, in a manner dependent on ZIP-1, a major transcription factor for IPR. Additionally, we found pathogen levels of the natural intracellular intestinal pathogen *Nematocida parisii* significantly reduced in *let-418* mutants, supporting the observation of increased IPR in this mutant. Altogether, these findings reveal a crucial role for LET-418 as a modulator of the IPR, aligning with its role in maintaining the balance between development and defense.

## Introduction

Chromatin is a highly dynamic structure, which regulates DNA access to the transcription machinery. It is regularly modified by large enzymatic complexes, including the widely conserved NuRD (Nucleosome Remodeling and Deacetylase) complex known to regulate developmental processes, as well as tissue homeostasis in many organisms (Xue *et al.* 1998; Shin and Mello 2003; Passannante *et al.* 2010; Reddy *et al.* 2010; Hu and Wade 2012; Hoffmann and Spengler 2019; Reid *et al.* 2022). Among the 7 unique subunits, each of them encoded by multiple gene paralogs, the NuRD complex relies on the CHD (chromodomain helicase DNA-binding) family of enzymes and histone deacetylases to regulate transcription (Reid *et al.* 2022). Classically it has been seen as a transcriptional repressor, however, there is evidence that the NuRD complex can also activate transcription (Bornelov *et al.* 2018; Zhang *et al.* 2018). The NuRD complex has several important roles, including being required for pluripotent cells to undergo lineage commitment in mice (Reynolds *et al.* 2012). The NuRD core component, CHD4/Mi-2β, is essential for stem cell homeostasis and lineage choice during skin development and mouse hematopoiesis (Kashiwagi *et al.* 2007; Yoshida *et al.* 2008). More recently, the NuRD complex was shown to be required for cardiac development and CHD4/Mi2β for proper skeletal muscle regeneration in mice (Wilczewski *et al.* 2018; Sreenivasan *et al.* 2021). Altogether, NuRD promotes cell fate transitions in a variety of different organisms and developmental contexts (Signolet and Hendrich 2015). In addition, NuRD subunits are frequently upregulated or mutated in cancers, indicating that this complex may regulate processes associated with tumorigenesis (Lai and Wade 2011).

In the nematode *Caenorhabditis elegans*, a NuRD-like complex is active in vulva formation (Solari and Ahringer 2000). The LET-418/Mi-2 NuRD member targets the HOX gene *lin-* 39 for repression, antagonizing the RAS signaling pathway that is required for vulval cell fate induction (Von Zelewsky *et al.* 2000; Guerry *et al.* 2007). LET-418/Mi-2 also plays a role in lifespan and stress resistance regulation. *let-418* mutant worms live longer and are more resistant to heat and oxidative stress (De Vaux *et al.* 2013). More recently, LET-418/Mi-2 and DCP-66/GATA2 NuRD members were shown to regulate multiple stress responses, including genotoxic stress and endoplasmic reticulum (ER) stress (Golden *et al.* 2022). In addition, LET-418 is required for germ cell fate maintenance. Together with the histone demethylase SPR-5/LSD1, it antagonizes the action of the COMPASS complex, maintaining proper H3K4 methylation levels in the germline and preventing germ cells to reprogram into somatic cells (KÄSER-PÉBERNARD *et al.* 2014). Beside the NuRD, LET-418/Mi-2 interacts with the conserved krüppel-type zinc-finger protein MEP-1, acting with the histone deacetylase HDA-1/HDAC1 to form a chromatin remodeling complex called the MEC complex. MEC complex members promote post-embryonic development, and they also repress the expression of germline genes in somatic cells (Unhavaithaya *et al.* 2002; Saudenova and Wicky 2018). A MEC complex, although lacking the histone deacetylase subunit, is also present in *Drosophila,* where it represses proneuronal genes during development (Kunert *et al.* 2009). Overall, LET-418/Mi-2 appears to function as a pro-developmental factor, which might prevent stress responses from being activated in order to maximize resource allocation to growth. Further investigations are required to understand how this factor balances growth and defense.

*C. elegans* can respond to a wide range of pathogens (Harding and Ewbank 2021). Intracellular pathogens, such as the Orsay virus and the fungus *Nematocida parisii,* which is part of the Microsporidia phylum, trigger a specific immune response called the intracellular pathogen response (IPR) (Troemel *et al.* 2008; Bakowski *et al.* 2014; Reddy *et al.* 2017; Tecle and Troemel 2022; L,aŽetić *et al.* 2023; Gang and Lazetic 2024). IPR is a transcriptional response that includes upregulation of many genes, including members of the *pals* gene family, named for the divergent ALS2CR12 protein sequence signature (Reddy *et al.* 2017; Reddy *et al.* 2019; Gang *et al.* 2022; Lazetic *et al.* 2022; Lazetic *et al.* 2023; Raman *et al.* 2024). This network of genes needs to be tightly controlled, since their deregulation leads to an imbalance between growth and defense. Thus, IPR positive regulators, such as *pals-20* and *pals-16* are specifically repressed during growth and development and cannot activate IPR effector genes, like *pals-5* and *pals-2* (Reddy *et al.* 2017; Lazetic *et al.* 2023). More recently, chromatin factors were also shown to play an important role in IPR regulation, as demonstrated by the function of MEP-1 and the NuRD member LIN-53 in IPR inhibition (Tecle *et al.* 2025). However, it remains to be determined how the IPR genes are specifically targeted by these upstream factors.

Here, we determined which genes are bound by LET-418/Mi-2 and MEP-1 in intestinal cells, as it was shown that intestinal tissue is the primary focus of action of LET-418/Mi-2. We found LET-418/Mi-2 and MEP-1 bound to genomic regions encompassing clusters of genes involved in innate immunity. Genes implicated in IPR are up-regulated in *let-418* mutants, which we show have increased resistance to microsporidia infection. Altogether these results indicate that LET-418/Mi-2 maintains the immune system in a repressed but inducible state, allowing normal growth and development.

## Results

### Identification of LET-418 and MEP-1 genomic binding sites using a DNA methylase technique

To better understand the function of LET-418 and MEP-1, we determined their genome binding sites specifically in the intestine, where we previously showed LET-418 to be required for post-embryonic development (Saudenova and Wicky 2018). To do so, we profiled the LET-418 and MEP-1 genome binding at the young adult stage using the DNA adenine methyltransferase identification (DamID) system (Gomez-Saldivar *et al.* 2021). Briefly, LET-418 and MEP-1 coding sequences are fused to Dam, which can methylate nearby nucleotides. Here, LET-418 and MEP-1 fusion constructs are preceded by a mCherry cassette that prevents their expression until it is excised out of the genome specifically by the CRE recombinase, whose expression is driven by the intestinal specific *elt-2* promoter (Figure 1A) (Fukushige *et al.* 1998). As a result, *let-418* or *mep-1* are expressed at very low level specifically in the intestine under the control of a non-induced *hsp-16.2* promoter. Following the experimental strategy depicted in Figure 1B, methylated fragments were PCR amplified and sequenced to generate the binding profile of LET-418 and MEP-1 (Sup Figure 1B, C). We used a nuclear diffusible GFP::Dam as negative control. Here we identified on average 5448 LET-418 peaks and 5245 MEP-1 peaks (Sup Figure 1D). Pairwise Pearson correlation (bin size 300 bp) between the replicates is high and MEP-1 and LET-418 binding profiles do not correlate with the control, indicating that MEP-1::Dam and LET-418::Dam fusions are introducing specific genome methylation (Figure 1C). Notably, we observe a high correlation (*r* > 0.5) between MEP-1 and LET-418 binding profiles (Figure 1C, 2A). The same trend is also observed with the Principal Component Analysis (PCA) (Sup Figure 1E). This finding suggests that LET-418 and MEP-1 share a significant number of binding sites. We analyzed the chromosome distribution of LET-418 and MEP-1 binding, but we did not find any specific regions that are enriched in LET-418 and MEP-1 binding sites, nor any bias in chromosome distribution (Figure 1D, E, Sup Figure 1F,G). Altogether, LET-418 and MEP-1 binding sites appear to be spread along all the chromosomes, but correlated with each other.

**Figure 1:**
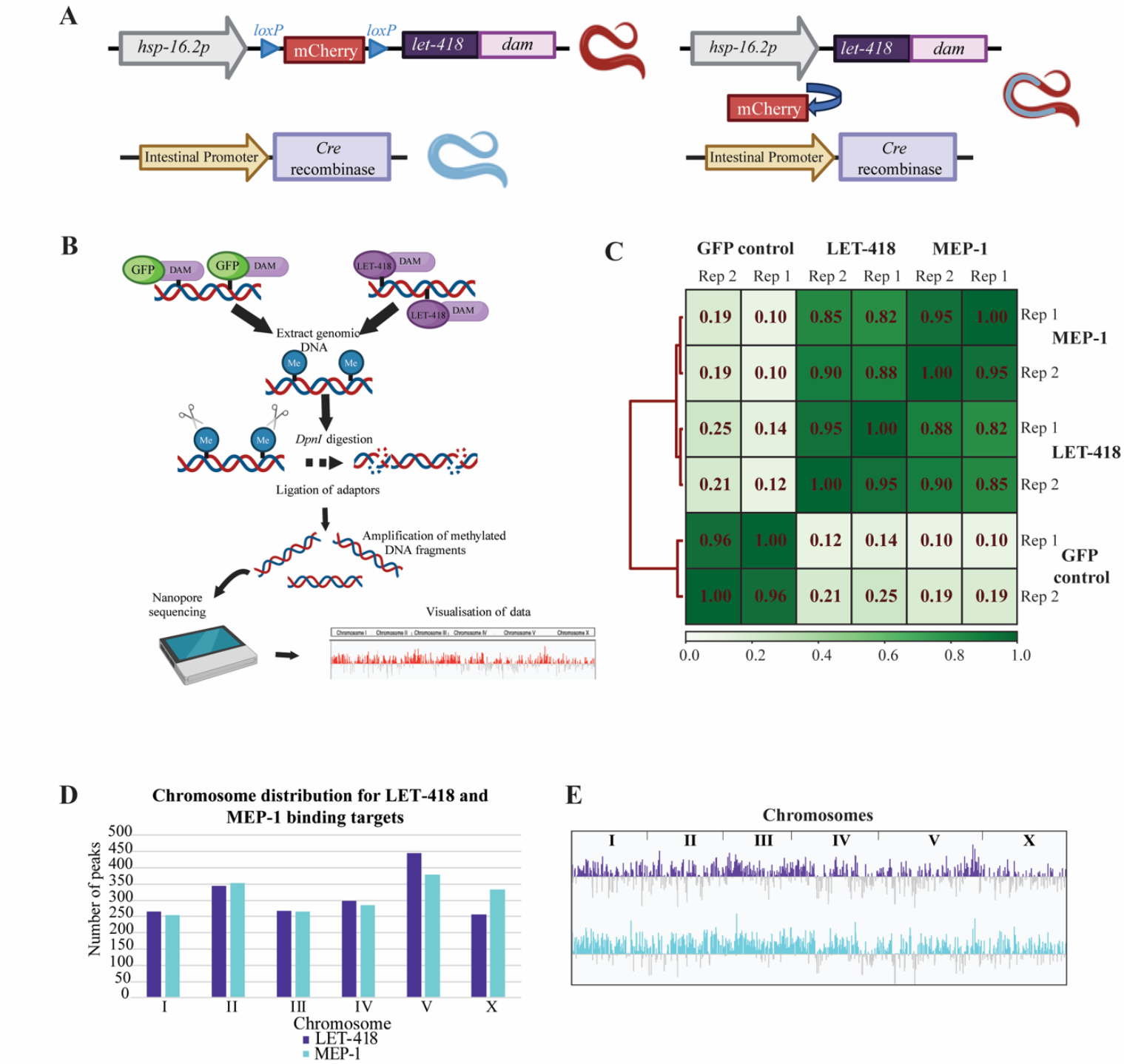
LET-418 and MEP-1 bind along the whole chromosome uniformly in intestinal cells. A) Tissue specific expression of LET-418::Dam using the Cre/lox recombination system.B) Experimental design of the DamID profiling. C) Pearson correlation between LET-418::Dam, MEP-1::Dam and NLS::GFP::Dam replicates. Red lines on the left indicate the relationship between each cluster. The length of the dendrogram branches shows the distance between clusters; the shortest being most related. The color bar represents the correlation between each sample with the darker shade indicating higher correlation. D) Chromosome distribution for LET-418 and MEP-1 target genes shows binding to all chromosomes with maximum binding on chromosome V. An average file of replicate 1 and 2 for both the proteins was used for the analysis. E) Binding profile of LET-418 and MEP-1 to all the chromosomes.

### LET-418 and MEP-1 share common target sites

LET-418 and MEP-1 binding profiles are highly correlated (*r* > 0.5) and taking a closer look at LET-418 and MEP-1 peak location revealed a very similar binding profile (Figure 2A). We generated an aggregation plot of LET-418 signal over MEP-1 peaks spanning a region of ±5 kb around peak signal midpoint, as well as the reciprocal analysis, and in both cases the findings are consistent with LET-418 and MEP-1 occupying common genomic loci (Figure 2B). This result agrees with LET-418 and MEP-1 being part of the same biochemical complex (Unhavaithaya *et al.* 2002; Passannante *et al.* 2010). Target gene identification (see material and methods) revealed that LET-418 binds to 1758 genes and MEP-1 to 1713 genes (Sup Table 1). Comparison of both genes lists shows that LET-418 and MEP-1 had 829 target genes in common (Figure 2C). Thus, only about half of LET-418 and MEP-1 targets appear to be unique to both proteins, indicating that MEP-1 and LET-418 have both common and unique biological functions. To identify the specific genomic elements that are bound by LET-418 and MEP-1, we assigned peaks to 2000 bp upstream and 2000 bp downstream of TSS (Transcriptional Start Site) and TES (Transcriptional End Site) respectively. LET-418 and MEP-1 both preferentially occupy an extended promoter region (Figure 2D). LET-418-bound DNA was previously shown in whole worms to be enriched with repetitive elements (Mcmurchy *et al.* 2017). We asked whether this is also the case in the intestine, but did not find any significant enrichment in repetitive elements (Sup Figure 2A, B). Interestingly, we observed that large families of genes organized in clusters, such as the F-box protein *fbxa* and the C-type lectin *clec* genes were heavily bound by LET-418 and MEP-1 (Sup Fig 2C, D). These clusters of genes, mostly containing stress response genes, are found in hypervariable regions across natural variants (Lee *et al.* 2021). Based on these findings, we wondered whether LET-418 and/or MEP-1 binding was enhanced in these hypervariable regions, however, we found no enrichment (Sup Figure 2E-G). In summary, LET-418 and MEP-1 appear to be enriched in non-coding chromatin, mostly 5’ end of genes, indicating a role in gene expression regulation.

**Figure 2:**
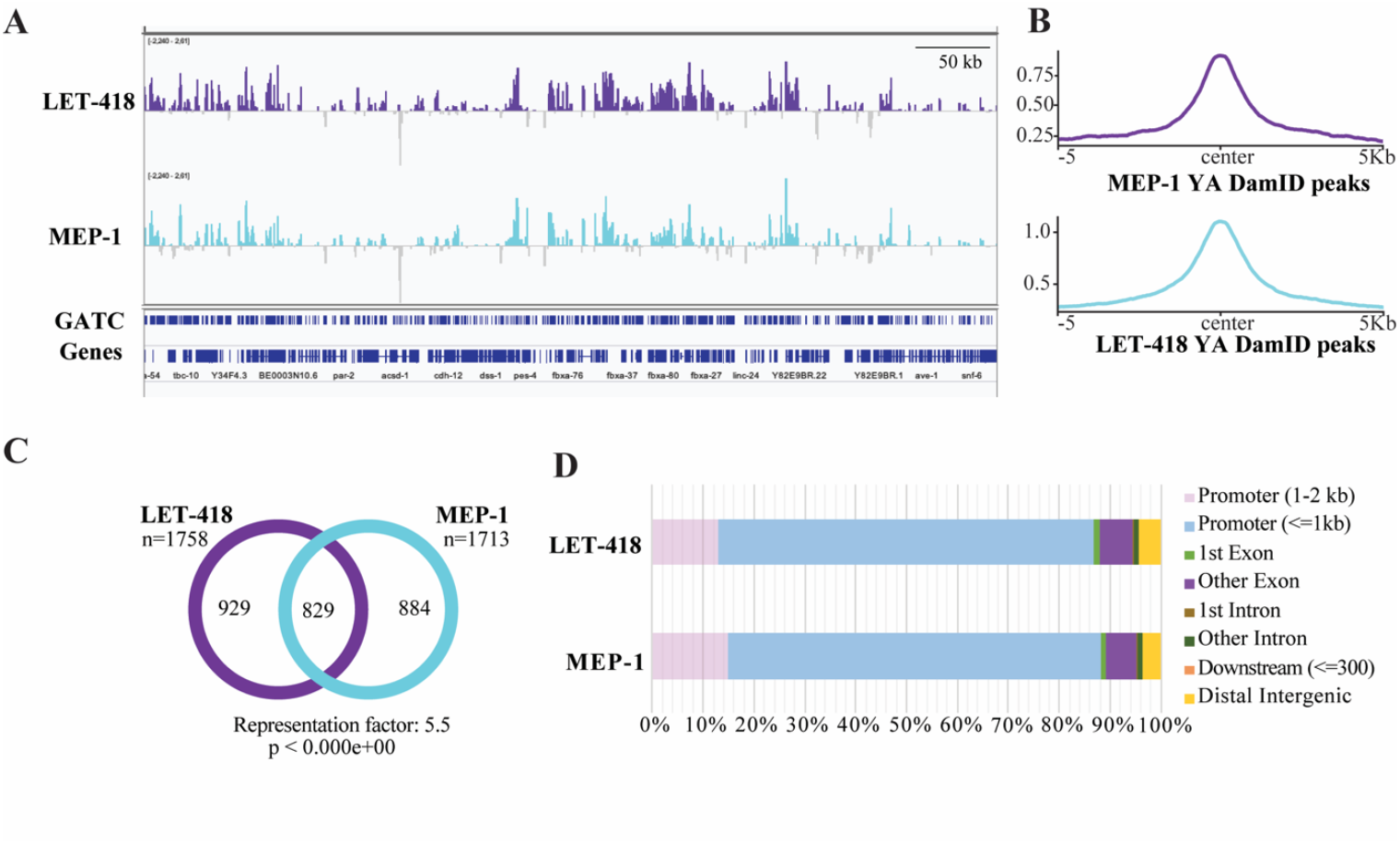
MEP-1 and LET-418 share half of their genomic targets. A) Representation on IGV browser of the LET-418 and MEP-1 signal profile in a selective region on Chromosome III (range -2.240-2.61). The *y* axes represent log_2_(LET-418/MEP-1::Dam/NLS::GFP::Dam) scores. B) Aggregation plots of LET-418::Dam signal over MEP-1::Dam peaks using with 5kb upstream and downstream of the center of the peaks and vice versa. YA = young adult stage.C) Venn diagram of the overlap between the target genes (average bedgraph file from both the replicates used to generate the gene list FDR<0.05) of LET-418 and MEP-1. The chart represents the number of genes identified for LET-418 and MEP-1. D) Bar chart representation of the genomic element distribution bound by LET-418 and MEP-1.

### LET-418 and MEP-1 binding regions are enriched in activating chromatin marks

To study the chromatin environment at LET-418 and MEP-1 binding sites, the LET-418 and MEP-1 binding profiles were compared with available profiles of young adult histone marks, which were processed from the ChIP-seq data from the ModERN resource (Kudron *et al.* 2018). Here we found that the most significant LET-418 and MEP-1 binding sites are highly enriched in histone marks associated with active transcription, such as H3K4me1/3, H3K36me2/3 and H3K79me3 (Figure 3A). In contrast, we found that modifications associated with negatively regulated genes, such as H3K9me2/3, are depleted from LET-418 and MEP-1 binding sites. However, H3K27me3, which also marks repressed genes, is enriched at MEP-1 binding sites, but not at LET-418 bound regions. LET-418 is often categorized as a heterochromatin protein, which prompted us to analyze the presence of the two heterochromatin proteins HPL-1/2 at LET-418 and MEP-1 binding sites (Ahringer and Gasser 2018). Using HPL-1/2 binding profiles generated by DamID specifically in the intestine and we observed that the two heterochromatin proteins were depleted from regions occupied by LET-418 (DE LA Cruz-Ruiz *et al.* 2023). HPL-1 showed depletion from MEP-1 bound regions while HPL-2 did not show any specific trend. The absence of HPL-1/2 from LET-418 or MEP-1 bound regions and vice versa, can be further visualized at specific genomic loci (Sup Figure 3A, B). In summary, LET-418 shows a distinct chromosome distribution compared to HPL-1/2 (Garrigues *et al.* 2015; Ahringer and Gasser 2018; DE LA Cruz-Ruiz *et al.* 2023). Using ChIP-seq data from the ModERN resource generated with whole worms at young adult stage, we further investigated the association of LET-418 and MEP-1 with additional heterochromatin associated proteins, namely the MBT domain protein LIN-61, the zinc-finger domain protein LIN-13, and the histone H3K9me1/2 methyltransferase MET-2 (Kudron *et al.* 2018). Out of this analysis, we found LIN-61 to be enriched at MEP-1 binding sites, while the other proteins tested showed no enrichment at MEP-1 and/or LET-418 bound regions (Figure 3C, D).

**Figure 3:**
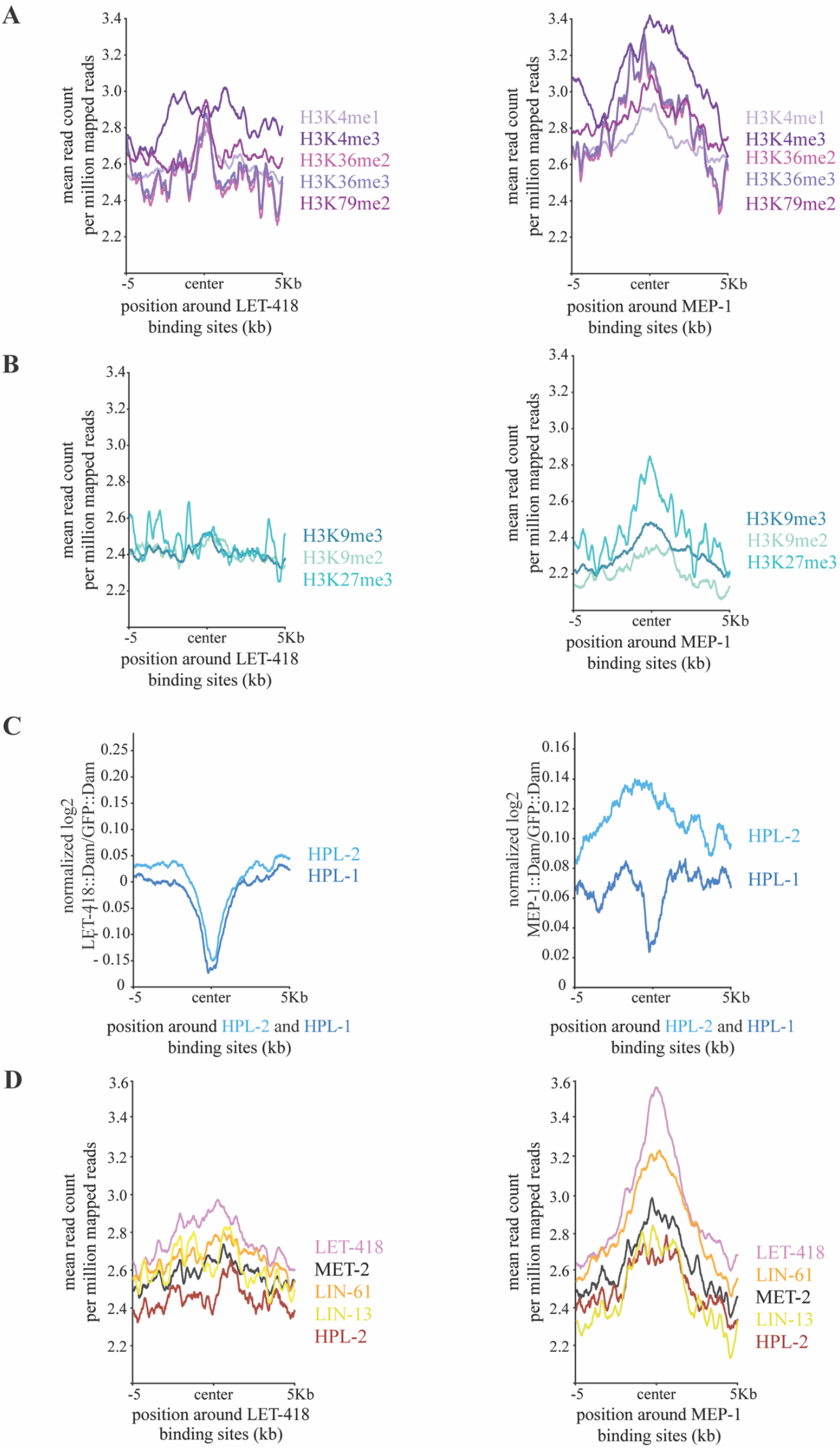
LET-418 and MEP-1 binding sites are enriched in histone marks associated with active transcription. Comparison of LET-418 and MEP-1 binding sites with known A) active (in violet) and B) repressive (in cyan) histone marks. C) Comparison of LET-418::Dam and MEP-1::Dam with HPL-1::Dam and HPL-2::Dam signals in the intestine. D) Comparison of LET-418 and MEP-1 binding sites with LIN-61, MET-2, LIN-13 and HPL-2 binding profiles. The reads were positioned around the center of the LET-418 and MEP-1 binding sites with 5000 bp regions upstream and downstream.

Altogether, our observations indicate that LET-418 and MEP-1 are associated with transcriptionally active chromatin and that they might function not to turn genes on and off, but rather to fine tune their expression. These observations agree with our previous findings in embryonic cells and with results obtained in stem cells (KÄser-pebernard *et al.* 2016; Bornelov *et al.* 2018). Interestingly, MEP-1 binding sites show an enrichment in H3K27me3 and LIN-61, an association that has not been previously detected.

### LET-418 and MEP-1 bind to target genes involved in intestinal functions and immune response

Tissue enrichment analysis revealed that the vast majority of genes bound by LET-418 and MEP-1 function in intestinal tissue (Figure 4A-C) (Angeles-Albores *et al.* 2016). For example, *zip-3* is highly enriched in both LET-418 and MEP-1 datasets, whereas *T19D12.4* is is bound by MEP-1. These genes are known to function in the intestine to regulate immune response (Sup Figure 4A). Interestingly, genes functioning in gonadal primordium are also enriched among LET-418 bound genes (Figure 4A). This finding is consistent with LET-418 being a repressor of germline genes in somatic cells (Unhavaithaya *et al.* 2002; Passannante *et al.* 2010). To dissect the shared and specific functions of LET-418 and MEP-1, we performed tissue enrichment analysis with genes that are only bound by either LET-418 or MEP-1. In these two gene lists, we also found strong enrichment in intestinal tissue (sup Figure 4B, C), however, functions in the muscular system are also associated with target genes unique to LET-418 (sup Figure 4A).

**Figure 4:**
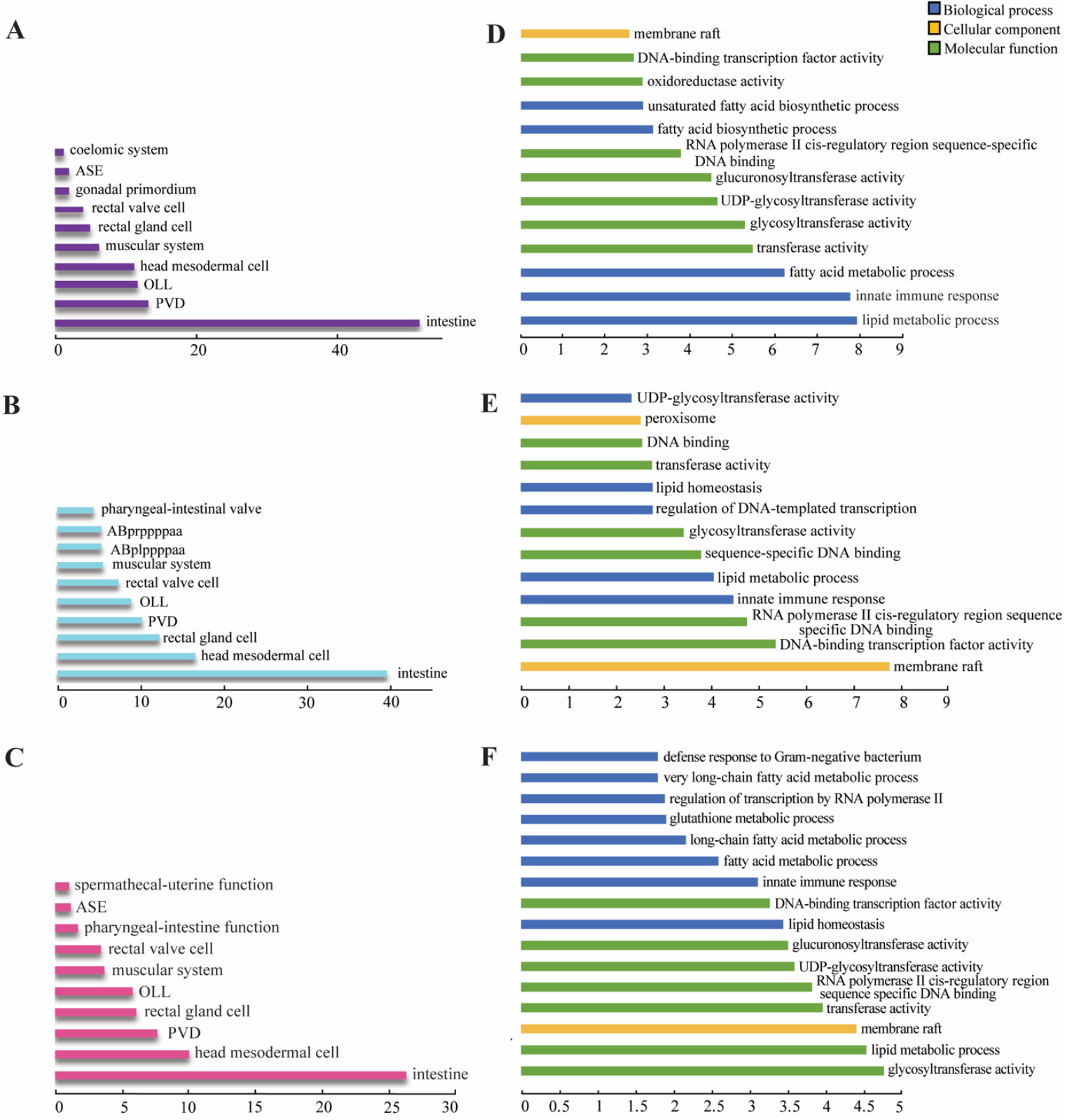
MEP-1 and LET-418 bind to genes functioning in the intestine in innate immunity and lipid metabolism. Tissue enrichment analysis for gene targets of A) LET-418 and B) MEP-1 as well as C) the common gene targets. q value threshold of 0.1. X axis is representing the –log10(q-value). DAVID GO term analysis for genes bound by D) LET-418 and E) MEP-1 as well as genes bound by F) both proteins using a background list of intestine specific genes (Kaletsky *et al.* 2018). X axis is representing the –log10(p-value).

To reveal functional involvement of LET-418 and MEP-1 in the intestine, we screened the gene target lists using the DAVID Gene Ontology (GO) term enrichment tool with an intestinal gene list as background (Kaletsky *et al.* 2018). We found that LET-418 and MEP-1 target genes are enriched in GO terms, such as innate immune response and lipid metabolism in the category of biological processes (Figure 4D, E). The same two biological processes are associated with target genes that are common to LET-418 and MEP-1 (Figure 4F). The term “membrane raft” emerged from the LET-418 and MEP-1 GO term enrichment analysis in the cellular component category, suggesting that they might regulate genes associated with membrane trafficking and cell signaling (Figure 4D-F). Next, we performed the same analysis on target genes unique to LET-418 or MEP-1. We found MEP-1 to be involved in sex-specific developmental processes (sup Figure 4D, E). The enrichment in genes involved in innate immunity in the list of LET-418 and MEP-1 common targets prompted us to look at their molecular nature in more details. As mentioned above, we observed that large family of genes, such as the *fbxa* and the *clec* genes were heavily bound by LET-418 and MEP-1 and involved in innate immunity (Sup Figure 2). LET-418 and MEP-1 also show binding to *pals* genes, which is a gene family expanded in *C. elegans.* The *pals* genes family includes members that are upregulated by intestinal intracellular infection, as well as genes that are not upregulated by infection, but which control expression of this transcriptional response. For example, we saw binding to *pals-20*, which regulates downstream *pals* genes that are upregulated as part of the intracellular pathogen response (IPR) (Sup Figure 4A left panel, Sup Table 5) (Lazetic *et al.* 2023). Altogether, these observations suggest that LET-418 and MEP-1 could play a role in modulating the immune response and more specifically the IPR.

### *let-418* loss-of-function mutants show upregulation of *pals* genes

To further investigate the link between LET-418 bound regions and gene regulation, we analyzed the transcriptome of *let-418(s1617)* loss-of-function mutant (Sup Figure 5A). Here we found that 2344 genes were upregulated while 1000 were downregulated (Figure 5A, Sup Table 6). Thus, most of the genes in the mutant strain were upregulated (2344/3344) suggesting a role for LET-418 in transcriptional repression. Interestingly, we found that 14 out of 39 *pals* genes are upregulated in the absence of LET-418 activity, and 12 of them are induced by *N. parisii* infection (Bakowski *et al.* 2014; Raman *et al.* 2024). Furthermore, GO analysis of the upregulated genes shows that immune response is among the processes regulated by LET-418 (Figure 5B). Additional GO terms found among up-regulated genes suggest a role of LET-418 in cell signaling. Down-regulated genes appear to be involved in nuclear functions and cell cycle (Figure 5C). Overlap between the differentially expressed genes and the LET-418 binding targets from the present DamID study revealed 260 genes in common out of which 160 genes are upregulated in the absence of LET-418 activity (Figure 5A). GO terms and tissue enrichment analysis indicate that LET-418 direct targets are functioning mainly in the intestine regulating immune response (sup Fig 5B-D).

**Figure 5:**
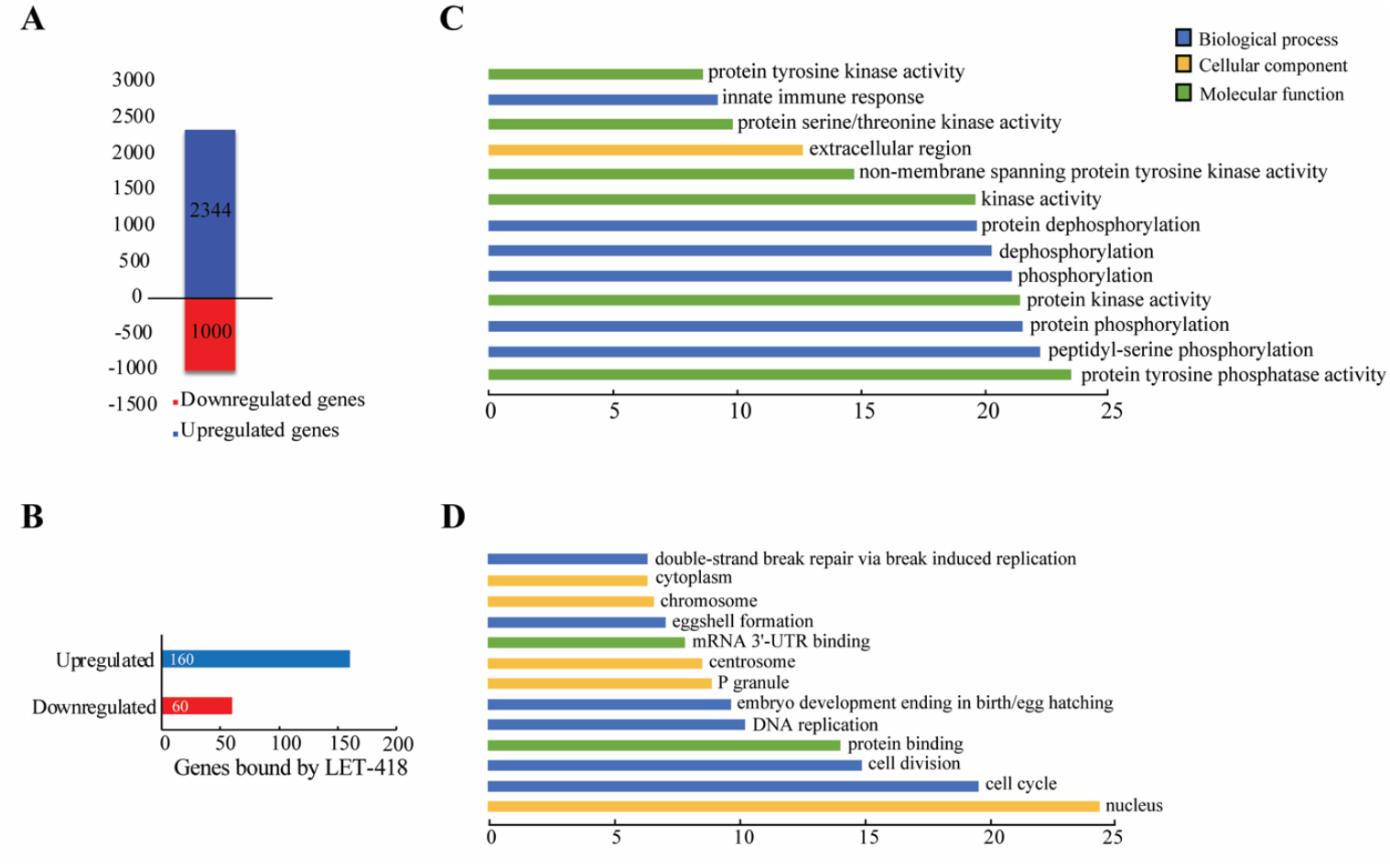
Absence of LET-418 leads to gene expression changes. A) Bar plot showing the number of upregulated and downregulated genes in *let-418* mutants. B) Bar plot representing the distribution of the deregulated genes in absence of LET-418, which are also bound by LET-418. DAVID GO term functional analysis for C) upregulated and D) downregulated genes in *let-418* mutants. X axis represents –log10(p-value).

### *let-418* mutants have increased resistance to intracellular pathogens

To further investigate the link between LET-418 and the IPR, we used the *pals-5p::*GFP transcriptional reporter, whose expression is upregulated in the absence of LET-418 activity as well as upon *N. parisii* infection, and is commonly used as a marker for IPR induction (Bakowski *et al.* 2014). *pals-5*p::GFP expression was induced in intestinal cells when *let-418* temperature sensitive mutants were grown at 25°C, (Figure 6A,C). In addition, we generated a single-copy translational GFP reporter for another IPR gene, *pals-2*, which was also found upregulated in the *let-418* mutant. Consistently, we observed a robust upregulation of *pals-2*::GFP expression in the intestine of *let-418* mutants grown at the restrictive temperature (Figure 6B, D). These results suggest that there is activation of the IPR in *let-418* mutants. The transcription factor ZIP-1 is known to function in the intestine downstream of all IPR-activating and regulatory pathways and it is required for the activation of *pals-5* expression upon infection by natural intracellular pathogens (Lazetic *et al.* 2023). To test if *pals-5* and *pals-2* upregulation is ZIP-1-dependent, we monitored expression of *pals-5* and *pals-2* reporters in *let-418(n3536) zip-1(RNAi)* worms. Here we observed that *pals-5* and *pals-2* induction was strongly suppressed upon *zip-1* knock-down, indicating that modulation of the IPR by LET-418 is *zip-1*-dependent (Figure 6A-D). Finally, to test whether IPR activation in *let-418* mutants increases resistance against microsporidia, we compared infection levels between *let-418* mutants and wild-type animals. We found a significantly lower number of sporoplasms (the early developmental stage of microsporidia) per animal in *let-418* mutants compared to the control strain, indicating that the loss of *let-418* provides immunity against microsporidia (Figure 6E). In summary, our results show that *let-418* mutants are more resistant to the intestinal intracellular pathogen *N. parisii*, most likely because of constitutive IPR induction.

**Figure 6:**
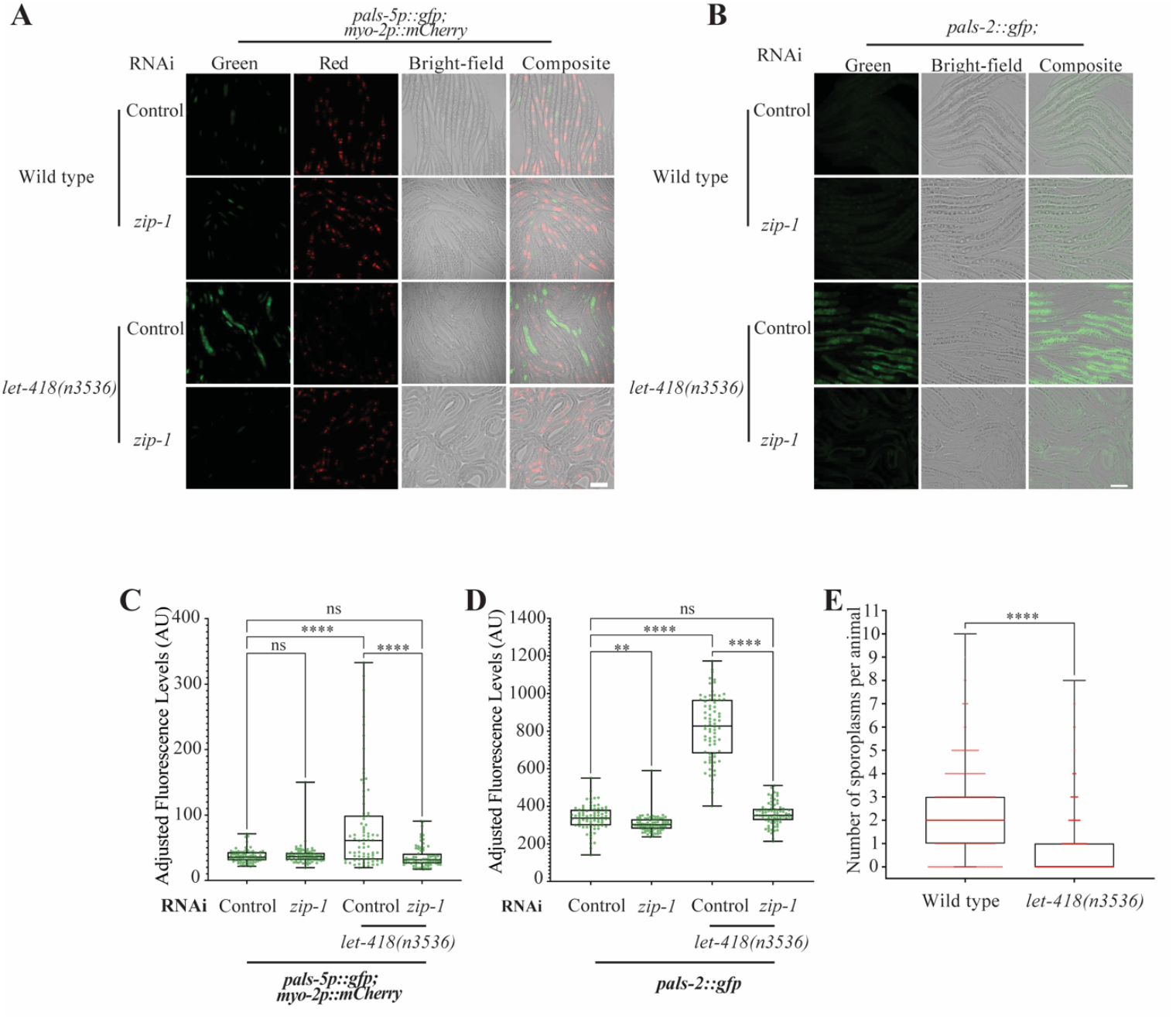
Expression of IPR reporters is induced in *let-418* mutants, which are more resistant to microsporidia infection. A) Representative images of *pals-5*p::GFP transcriptional reporter expression in wild-type worms and *let-418* mutants treated for 6 hours with control RNAi or *zip-1* RNAi. *myo-2*p::mCherry in red was used as a co-injection marker B) Representative images of *pals-2*::GFP translational reporter expression in wild type worms and *let-418* mutants treated for 23 hours with control or *zip-1* RNAi. Scale bar is 50 µM. C, D) Quantification of the fluorescence was performed for the indicated strains and represented as a box-and-whiskers plot. The line in the box represents the median value. Boxes bound indicate 25th and 75th percentiles, and whiskers extend from the box bounds to the minimum and maximum values. A Kruskal-Wallis test was applied to calculate the p-values E) Wild type worms and *let-418* mutants were infected by *N. parisii* and infection level was measured by scoring the number of sporoplasms per animals (y axis) for each genotype (x axis). A Kolmogorov-Smirnov test was used to calculate p-values; **** p < 0.0001; ** 0.001 ≤ p < 0.01; ns indicates nonsignificant difference (p > 0.05).

## Discussion

Using DamID profiling, we found that LET-418 and MEP-1 bind to common genomic regions distributed uniformly across all chromosomes. Chromatin associated with LET-418 and MEP-1 binding sites appears to be enriched in histone modifications that are linked to active transcription, and functional analysis of LET-418 and MEP-1 target genes reveal an enrichment in genes involved in innate immunity. We observed that *let-418* mutants had upregulation of genes associated with the IPR, a response to intracellular intestinal pathogens. Accordingly, we also found that *let-418* mutants have increased resistance to an intestinal intracellular pathogen *N. parisii*. Based on these results, we propose a model whereby LET-418 and MEP-1 keep immune response gene expression at a low transcriptional level, but in a chromatin state that is responsive. Upon infection, chromatin is remodeled, and the expression of immune genes can be induced by the ZIP-1 transcription factor (Figure 7).

**Figure 7:**
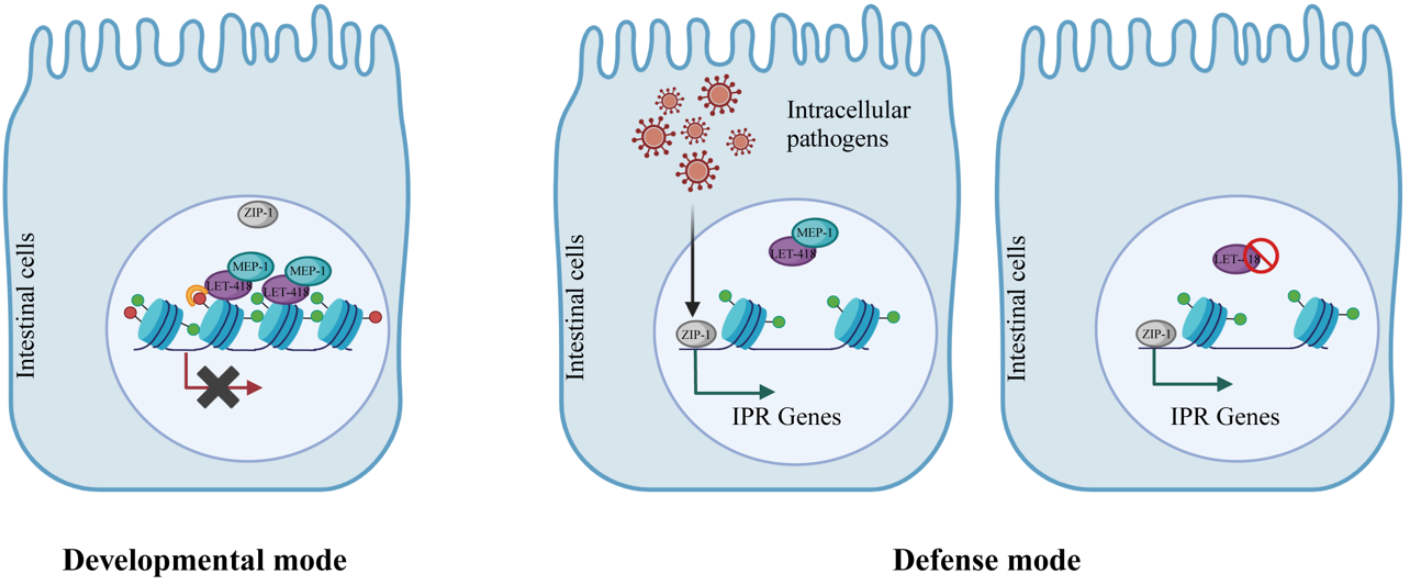
Chromatin is kept responsive by LET-418 and MEP-1. It is re-shaped upon intracellular pathogen infection via the action of ZIP-1 (for details: see text).

LET-418 and MEP-1 share many binding sites, suggesting that they are acting together in a complex. In agreement with this model, *let-418* and *mep-1* mutants also exhibit developmental defects in common. Absence of *let-418* or *mep-1* maternal activity leads to developmental arrest at the first larval stage, associated with ectopic expression of germline genes in the soma (Von Zelewsky *et al.* 2000; Unhavaithaya *et al.* 2002). At L1 stage, deregulated genes in *let-418* and *mep-1* mutants exhibit more than 80% overlap (Passannante *et al.* 2010). Our DamID profiling experiments performed at adult stage show that around 50% of the target genes are shared by both proteins, pointing at shared function performed most likely by a MEC complex, however, both proteins also bind to unique target sites. LET-418 is also a member of the NuRD complex that is involved in many biological functions, including development and stress response (Passannante *et al.* 2010; Zhu *et al.* 2020; Golden *et al.* 2022).

LET-418 is often associated with heterochromatin proteins, functioning to repress transcription (Mcmurchy *et al.* 2017; Ahringer and Gasser 2018). However, we found that its binding pattern differs from the heterochromatin proteins HPL-1/2. In the intestine, our data shows that LET-418 occupies sites that are depleted in HPL-1/2, suggesting that LET-418 is not associated with heterochromatic regions (DE LA Cruz-Ruiz *et al.* 2023). In line with these observations, LET-418 bound regions do not appear to be enriched in repetitive elements, and are rather enriched in histone modifications related to active transcription. We observed a similar association in whole worms at the embryonic stage (KÄser-pebernard *et al.* 2016; Hou *et al.* 2023). However, an association with H3K9me2 was also reported (Mcmurchy *et al.* 2017). Altogether, this suggests a role for LET-418 and MEP-1 in the modulation of the level of transcription. In this model, LET-418 and MEP-1 would keep transcriptional levels below a certain threshold, rather than functioning as an on/off switch. In mammals, binding of the LET-418 homolog CHD4 correlates with methylated H3K4, an association that has been proposed to make regulatory sequences responsive to internal and external cues (Bornelov *et al.* 2018). Our data indicates that LET-418 and MEP-1 b2ind to regions near genes that mainly function in the intestine. LET-418 is required in the intestine to promote post-embryonic development, suggesting that LET-418 targeted genes in the intestine are important for growth and development (Saudenova and Wicky 2018). However, functional analysis revealed that target genes are enriched in innate immunity-associated genes, a signature that is also seen in transcriptomic data (own data and (Golden *et al.* 2022)). Altogether, these results suggest that LET-418 and MEP-1 are repressing defense genes but maintaining chromatin in a responsive state. This interpretation is in agreement with the observation that misregulation of defense mechanisms leads to growth defects and further highlights the necessity of regulating defense genes to allow development (Lazetic *et al.* 2023). LET-418 and MEP-1 could be counteracted by immune response regulators such as the ZIP-1 transcription factor. Still in *C. elegans*, a similar type of regulation was demonstrated for the THAP (Thanatos associated protein)-domain protein LIN-15B and members of the DRM (DREAM) complex including LIN-35, which were shown to negatively regulate the IPR (Tecle *et al.* 2025). Interestingly, LIN-15B and DRM complex members are all part of the Synthetic Multivulva (SynMuv) family of developmental genes, whose other members, including MEP-1 and the NuRD complex member LIN-53 were shown to upregulate the transcriptional IPR reporter *pals-5*p::GFP (Tecle *et al.* 2025).

Activation of IPR in the *let-418* mutants could also be viewed as an indirect consequence of defects associated with lack of LET-418 activity. It is possible that massive gene deregulation caused by the absence of LET-418 could trigger homeostasis-altering molecular processes (HAMPs), which in turn activates innate immunity and stress resistance, leading to developmental delay, and pathogen and stress resistance (De Vaux *et al.* 2013; Liston and Masters 2017). Alternatively, DNA damage in the germ cells generated by the failure to repair double-strand brakes during meiotic recombination in the *let-418* mutant could induce innate immunity and systemic stress resistance, a mechanism that was proposed to reinforce somatic tissue maintenance (Ermolaeva *et al.* 2013; Turcotte *et al.* 2018).

In summary, at the organismal level, LET-418 and MEP-1 positively regulate growth and developmental processes, while limiting defense mechanisms. LET-418 function is conserved, since the mammalian homolog CHD4/Mi-2β,is also driving development, however, its potential role in defense mechanisms highlighted here using *C. elegans* remains to be determined.

## Materials and Methods

### *C. elegans* maintenance and growth of strains

*C. elegans* strains were maintained as per standard protocols on a lawn of OP50 *Escherichia coli* on nematode growth medium (NGM) plates at 20°C. For the DamID protocol, *C. elegans* strains were grown on a lawn of GM48 (-dam) *E. coli* bacteria on NGM plates at 20°C for at least two generations. After synchronization, around 1500 L1 larvae were seeded on three 100 mm plates. The larvae were grown until the young adult stage and were collected after washing 10-12 times with M9 media in aliquots of 30-50 µl. After the removal of excess liquid, samples (approximately 4500 worms per sample) were snap frozen and stored at -80°C until further use. The list of strains used in the study is mentioned in Sup Table 2.

### Molecular cloning and MosSCI insertion

Tissue-specific plasmids required for DamID experiments were generated using the Gibson Assembly technique (Gibson *et al.* 2009). Generation of the plasmid for the *pals-2* translational reporter strain used Gateway (Invitrogen) cloning and insertion of GFP fragment using restriction digest. Integration of the plasmids into the genome was done using the MosSCI insertion (Frokjaer-Jensen *et al.* 2008) and were inserted on Chromosome II or IV as single copy plasmids. The list of plasmids and primers used in this study are mentioned in Sup Tables 3 and 4.

### DamID seq protocol

All the steps for the DamID experiments including the molecular cloning to the multiplexed library preparation were followed according to (Gomez-Saldivar *et al.* 2021) and graphically described in Figure 1B. We confirmed excision of the mCherry cassette by PCR (sup Figure 1A). DNA extraction from frozen worm pellets was performed by using the DNeasy Blood and Tissue Kit (#69504; Qiagen, Valencia, CA), and further concentrated using ethanol precipitation. All the experiments were conducted using two biological replicates for each strain. 500 ng of DNA was used for every sample for digestion with *DpnI* followed by ligation of the adaptors and PCR amplification to generate DNA amplicons of each sample. For the intestinal DamID 18-22 PCR cycles were performed (sup Figure 1B, representative replicate 1). DNA amplicon purifications were performed after every step from the PCR amplification onwards using magnetic beads and DNA quantification was done using QUBIT. Library preparation included end repair, barcoding and pooling of the libraries to be loaded on the MinION mk1c Nanopore Sequencer. The run lasted for 72 hours yielding approximately 10 million reads, around one million reads per library (sup Figure 1C).

### DamID data processing

The raw data obtained from the Oxford Nanopore Sequencing runs are in the form of a compressed fast5 file. This file is then processed in a number of steps including base-calling, quality check, demultiplexing, read mapping, DamID read filtering and finally DamID signal determination. The detailed steps for the bioinformatic analysis have been mentioned in (Gomez-Saldivar *et al.* 2021). Normalization for the protein of interest (POI) for GATC accessibility was calculated as the log2 ratio between the POI:: Dam signal versus the GFP::Dam (freely moving). The pipeline used to determine the DamID signal profile was downloaded from https://github.com/owenjm/damidseq_pipeline. The input for this pipeline is the bam file generated from raw data processing and the output is a bedGraph file which is used to visualize peaks or the binding regions of the protein of interest on the IGV browser. The number of peaks for each replicate is mentioned in sup Figure 1D. The normalized log2 POI::Dam/GFP::Dam bedGraph file is then used for the further downstream analysis.

### Peak calling and assignment of peaks

BedGraph files for both the replicates from the previous step were used as an input to generate an average of the bedgraph files using the average tracks perl script available at https://github.com/owenjm/damid_misc. The average file was used to find significant peaks (false discovery rate (FDR < 0.05) with the output being a GFF file, using find_peaks Perl script (downloaded from https://github.com/owenjm/find_peaks). Gene annotation of the statistically significant peaks (FDR< 0.05) was performed using the R package ChIPpeakAnno. Gene annotation was done using the reference genome *C. elegans* WBcel235 using the R package ensembldb. The R script is available on request. The average bedGraph files for each POI were used for visualization on IGV browser, generation of aggregation plots, analysis of genomic element and repeat element distribution, as well as hypervariable region enrichment. Venn diagrams for the comparison of overlaps between target genes of various datasets were performed using the online tool https://bioinformatics.psb.ugent.be/webtools/Venn/ and the statistical significance of the overlaps between the datasets was calculated using http://nemates.org/MA/progs/overlap_stats.html.

### Pearson’s correlation

The Pearson’s correlation and principal component analysis was done using the online platform https://usegalaxy.org. The Pearson’s correlation was done using the multiBamSummary tool with a bin size of 300 bp. To plot the heatmap of the correlation between the samples, plotCorrelation tool was used on the multiBamSummary matrix.

### Aggregation plots and genomic element distribution

The usegalaxy.org platform was used to generate aggregation plots and genomic element distribution. Aggregation plots comparing DamID data with other DamID data or ChIP-seq data were done using computeMatrix. The regions to be plotted were provided as a gff/bed file while the file to be scored were in the form of bedgraph/bigwig files. The reference point for the plotting was the center of the GFF/bed file and genomic regions 5000 bp upstream and downstream were considered. A bin size of 10 bp was used. The output file was used in the plotProfile tool to generate the aggregation plots and to observe the signal localization. For the representation of genomic element distribution for LET-418 and MEP-1, ChIPseeker tool was used. The hypervariable regions co-ordinates to look at the LET-418 and MEP-1 occupancy were obtained from (Lee *et al.* 2021). Chromatin immunoprecipitation data were obtained from the following studies: H3K4me1,H3K4me3, H3K36me3 (Janes *et al.* 2018), H3K9me2, H3K9me3, HPL-2, LIN-13, MET-2, LIN-61, LET-418 (Mcmurchy *et al.* 2017); H3K36me2, H3K27me3 (Zaghet *et al.* 2021) and H3K79me3 from modENCODE Consortium (https://www.encodeproject.org/) (Kudron *et al.* 2018). The NCBI SRR accession numbers of all the files are mentioned in Sup Table 7.

### Tissue enrichment and gene ontology

Tissue enrichment analysis for the datasets was performed using the online tool available on Wormbase (https://wormbase.org/tools/enrichment/tea/tea.cgi) (Angeles-Albores *et al.* 2016). Gene ontology analysis was done using the DAVID Functional Annotation Bioinformatics Microarray Analysis (https://david.ncifcrf.gov/tools.jsp) (Sherman *et al.* 2022).

### Repeat element distribution

The reference repetitive elements list for *C. elegans* (WBcel235/ce11) was downloaded from the UCSC genome browser using the table browser tool. Group selected was variations and repeats and downloaded as a bed file. This file was edited to exclude simple repeats and low complexity regions. Bedtools intersect tool was used to get the intersection between the reference file and the DamID datasets. Bedtools fisher tool was used to determine the significance of this overlap.

### Microsporidia infections

*N. parisii* spores were prepared as previously described in (Balla *et al.* 2015). 1.26 million spores were combined with 10 µl 10x OP50 *E. coli*, 1,200 synchronized L1-stage *C. elegans*, and M9 buffer (total volume 300 µl). This mixture was then plated on 6 cm NGM plates, allowed to dry for 30 min at room temperature, and then incubated at 25 °C for 3 hours. Following incubation, the worms were collected using M9 buffer with 0.1% Tween 20, washed with PBS containing 0.1% Tween 20 (PBST), and fixed in 4% paraformaldehyde for 30 minutes. The fixed samples were then stained overnight at 46 °C using FISH probes conjugated with the red Cal Fluor 610 fluorophore, targeting ribosomal RNA (Rivera *et al.* 2022). Analysis was performed using a Zeiss Axio Imager.M1 compound microscope. The data were plotted and statistically analyzed with GraphPad Prism software.

### RNAi assays

RNA interference (RNAi) assays were performed using the feeding method (Timmons *et al.* 2001). Overnight cultures of HT115 *E. coli* were spread on RNAi plates (NGM plates supplemented with 5 mM IPTG and 1 mM carbenicillin) and incubated at room temperature in the dark for at least three days. RNAi clones were sourced from the Ahringer RNAi library, with the vector plasmid L4440 serving as a negative control. L4-stage animals were transferred to RNAi plates and maintained at 20 °C for four days. Gravid adults were then bleached to get eggs, that were incubated in M9 medium overnight to hatch into starved L1 larvae. Subsequently, 1,000 synchronized L1 larvae were transferred to RNAi plates. The plates were dried for 30 min at room temperature, then incubated at 20 °C for one hour, followed by incubation at 25 °C for 6 h (for animals with the *pals-5p::gfp* reporter) or 23 h (for animals with the *pals-2::gfp* reporter). Animals were anesthetized using 20 µM levamisole and mounted on agarose pads on glass slides for imaging. Imaging was conducted using a Zeiss Axio Imager.M2 compound microscope. The mean gray value (calculated as the ratio of integrated density to analyzed area) was determined for each animal and normalized against background fluorescence in FIJI program. The data were plotted and statistically analyzed with GraphPad Prism software

### RNA isolation and Transcriptomic analysis

Wild type worms and *let-418(s1617)/tmC16* mutant worms were synchronised to allow collection of around 1500 young adults. Three biological replicates of each strain were collected. RNA extraction was performed using TRIzol reagent (Invitrogen, Carlsbad CA, and then purified using the PureLink RNA Mini Kit (Invitrogen, Carlsbad CA,USA). The RNA samples were further processed and sequenced at the Next Generation Sequencing (NGS) Platform in Bern (https://www.ngs.unibe.ch). Data received in fasta format from NGS platform Bern was uploaded on https://useglaxy.org online platform for further analysis. The pipeline for analysis was followed as mentioned in (Rodriguez-Crespo *et al.* 2022).

## Acknowledgements

The authors would like to thank the *Caenorhabditis* Genetics Center (CGC; cbs.umn.edu/cgc/home) and *C. elegans* Gene Knockout Consortium by S. Mitani at the Tokyo Women’s Medical University School of Medicine (Tokyo, Japan) for providing strains required for the experiments.

## Data Availability

Raw files and processed files from the DamID experiment and RNA sequencing are available on ArrayExpress database at EMBL-EBI (www.ebi.ac.uk/arrayexpress) with accession numbers E-MTAB-14824 and E-MTAB-14522 respectively. Details for the source of ChIP-seq data are mentioned in the Supplementary Table 7. Strains and plasmids are available on request.

## Funding

This work was supported by IZCOZ0_198093 (linked to COST Action CA18127 International Nucleome Consortium) and SNSF (Swiss National Science Foundation) Grants 31003A_179395 to CW.

## Notes

https://www.ebi.ac.uk/biostudies/arrayexpress/studies/E-MTAB-14824

https://www.ebi.ac.uk/biostudies/arrayexpress/studies/E-MTAB-14522

### Competing Interest Statement

The authors have declared no competing interest.

### Summary of Updates

Nomenclature in the title edited to italics.

## Bibliography

Ahringer, J., and S. M. Gasser, 2018 Repressive Chromatin in Caenorhabditis elegans: Establishment, Composition, and Function. Genetics 208: 491–511.

Angeles-Albores, D., N. L. RY, J. Chan and P. W. Sternberg, 2016 Tissue enrichment analysis for C. elegans genomics. BMC Bioinformatics 17: 366.

Bakowski, M. A., C. A. Desjardins, M. G. Smelkinson, T. L. Dunbar, I. F. Lopez-Moyado et al., 2014 Ubiquitin-mediated response to microsporidia and virus infection in C. elegans. PLoS Pathog 10: e1004200.

Balla, K. M., E. C. Andersen, L. Kruglyak and E. R. Troemel, 2015 A wild C. elegans strain has enhanced epithelial immunity to a natural microsporidian parasite. PLoS Pathog 11: e1004583.

Bornelov, S., N. Reynolds, M. Xenophontos, S. Gharbi, E. Johnstone et al., 2018 The Nucleosome Remodeling and Deacetylation Complex Modulates Chromatin Structure at Sites of Active Transcription to Fine-Tune Gene Expression. Mol Cell 71: 56–72 e54.

de la Cruz-Ruiz, P. M. J. Rodriguez-Palero, P. Askjaer and M. Artal-Sanz, 2023 Tissue-specific chromatin-binding patterns of Caenorhabditis elegans heterochromatin proteins HPL-1 and HPL-2 reveal differential roles in the regulation of gene expression. Genetics 224: iyad081.

De Vaux, V., C. Pfefferli, M. Passannante, K. Belhaj, A. von Essen et al., 2013 The Caenorhabditis elegans LET-418/Mi2 plays a conserved role in lifespan regulation. Aging Cell 12: 1012–1020.

Ermolaeva, M. A., A. Segref, A. Dakhovnik, H. L. Ou, J. I. Schneider et al., 2013 DNA damage in germ cells induces an innate immune response that triggers systemic stress resistance. Nature 501: 416–420.

Frokjaer-Jensen, C., M. W. Davis, C. E. Hopkins, B. J. Newman, J. M. Thummel et al., 2008 Single-copy insertion of transgenes in Caenorhabditis elegans. Nat Genet 40: 1375–1383.

Fukushige, T., M. G. Hawkins and J. D. McGhee, 1998 The GATA-Factor elt-2 Is Essential for Formation of the Caenorhabditis elegans Intestine. Dev Biol 198: 286–302.

Gang, S. S., M. Grover, K. C. Reddy, D. Raman, Y. T. Chang et al., 2022 A pals-25 gain-of-function allele triggers systemic resistance against natural pathogens of C. elegans. PLoS Genet 18: e1010314.

Gang, S. S., and V. Lazetic, 2024 Microsporidia: Pervasive natural pathogens of Caenorhabditis elegans and related nematodes. J Eukaryot Microbiol 71: e13027.

Garrigues, J. M., S. Sidoli, B. A. Garcia and S. Strome, 2015 Defining heterochromatin in C. elegans through genome-wide analysis of the heterochromatin protein 1 homolog HPL-2. Genome Res 25: 76–88.

Gibson, D. G., L. Young, R. Y. Chuang, J. C. Venter, C. A. Hutchison, 3rd et al., 2009 Enzymatic assembly of DNA molecules up to several hundred kilobases. Nat Methods 6: 343–345.

Golden, N. L., M. K. Foley, K. S. Kim Guisbert and E. Guisbert, 2022 Divergent regulatory roles of NuRD chromatin remodeling complex subunits GATAD2 and CHD4 in Caenorhabditis elegans. Genetics 221: iyac046.

Gomez-Saldivar, G., D. A. Glauser and P. Meister, 2021 Tissue-specific DamID protocol using nanopore sequencing. J Biol Methods 8: e152.

Guerry, F., C. O. Marti, Y. Zhang, P. S. Moroni, E. Jaquiery et al., 2007 The Mi-2 nucleosome-remodeling protein LET-418 is targeted via LIN-1/ETS to the promoter of lin-39/Hox during vulval development in C. elegans. Dev Biol 306: 469–479.

Harding, B. W., and J. J. Ewbank, 2021 An integrated view of innate immune mechanisms in C. elegans. Biochem Soc Trans 49: 2307–2317.

Hoffmann, A., and D. Spengler, 2019 Chromatin Remodeling Complex NuRD in Neurodevelopment and Neurodevelopmental Disorders. Front Genet 10: 682.

Hou, X., M. Xu, C. Zhu, J. Gao, M. Li et al., 2023 Systematic characterization of chromodomain proteins reveals an H3K9me1/2 reader regulating aging in C. elegans. Nat Commun 14: 1254.

Hu, G., and P. A. Wade, 2012 NuRD and pluripotency: a complex balancing act. Cell Stem Cell 10: 497–503.

Janes, J., Y. Dong, M. Schoof, J. Serizay, A. Appert et al., 2018 Chromatin accessibility dynamics across C. elegans development and ageing. Elife 7: e37344.

Kaletsky, R., V. Yao, A. Williams, A. M. Runnels, A. Tadych et al., 2018 Transcriptome analysis of adult Caenorhabditis elegans cells reveals tissue-specific gene and isoform expression. PLoS Genet 14: e1007559.

Käser-Pébernard, S., F. Müller and C. Wicky, 2014 LET-418/Mi2 and SPR-5/LSD1 Cooperatively Prevent Somatic Reprogramming of C. elegans Germline Stem Cells. Stem Cell Reports 2: 547–559.

Käser-Pebernard, S., C. Pfefferli, C. Aschinger and C. Wicky, 2016 Fine-tuning of chromatin composition and Polycomb recruitment by two Mi2 homologues during C. elegans early embryonic development. Epigenetics Chromatin 9: 39.

Kashiwagi, M., B. A. Morgan and K. Georgopoulos, 2007 The chromatin remodeler Mi-2b is required for establishment of the basal epidermis and normal differentiation of its progeny. Development 134: 1571–1582.

Kudron, M. M., A. Victorsen, L. Gevirtzman, L. W. Hillier, W. W. Fisher et al., 2018 The ModERN Resource: Genome-Wide Binding Profiles for Hundreds of Drosophila and Caenorhabditis elegans Transcription Factors. Genetics 208: 937–949.

Kunert, N., E. Wagner, M. Murawska, H. Klinker, E. Kremmer et al., 2009 dMec: a novel Mi-2 chromatin remodelling complex involved in transcriptional repression. EMBO J.

Lai, A. Y., and P. A. Wade, 2011 Cancer biology and NuRD: a multifaceted chromatin remodelling complex. Nat Rev Cancer 11: 588–596.

Lažetić, V., L. E. Batachari, A. B. Russell and E. R. Troemel, 2023 Similarities in the induction of the intracellular pathogen response in Caenorhabditis elegans and the type I interferon response in mammals. BioEssays 45: 202300097.

Lazetic, V., M. J. Blanchard, T. Bui and E. R. Troemel, 2023 Multiple pals gene modules control a balance between immunity and development in Caenorhabditis elegans. PLoS Pathog 19: e1011120.

Lazetic, V., F. Wu, L. B. Cohen, K. C. Reddy, Y. T. Chang et al., 2022 The transcription factor ZIP-1 promotes resistance to intracellular infection in Caenorhabditis elegans. Nat Commun 13: 17.

Lee, D., S. Zdraljevic, L. Stevens, Y. Wang, R. E. Tanny et al., 2021 Balancing selection maintains hyper-divergent haplotypes in Caenorhabditis elegans. Nat Ecol Evol 5: 794–807.

Liston, A., and S. L. Masters, 2017 Homeostasis-altering molecular processes as mechanisms of inflammasome activation. Nat Rev Immunol 17: 208–214.

McMurchy, A. N., P. Stempor, T. Gaarenstroom, B. Wysolmerski, Y. Dong et al., 2017 A team of heterochromatin factors collaborates with small RNA pathways to combat repetitive elements and germline stress. Elife 6: 21666.

Passannante, M., C. O. Marti, C. Pfefferli, P. S. Moroni, S. Kaeser-Pebernard et al., 2010 Different Mi-2 Complexes for Various Developmental Functions in Caenorhabditis elegans. PLoS One 5: e13681.

Raman, D., N. Wernet, S. S. Gang and E. R. Troemel, 2024 PALS-14 promotes resistance to Nematocida parisii infection in Caenorhabditis elegans. microPublication Biology: 10.17912/micropub.biology.001325.

Reddy, B. A., P. K. Bajpe, A. Bassett, Y. M. Moshkin, E. Kozhevnikova et al., 2010 Drosophila Transcription Factor Tramtrack69 Binds MEP1 to Recruit the Chromatin Remodeler NuRD. Mol Cell Biol.

Reddy, K. C., T. Dror, J. N. Sowa, J. Panek, K. Chen et al., 2017 An Intracellular Pathogen Response Pathway Promotes Proteostasis in C. elegans. Curr Biol 27: 3544–3553 e3545.

Reddy, K. C., T. Dror, R. S. Underwood, G. A. Osman, C. R. Elder et al., 2019 Antagonistic paralogs control a switch between growth and pathogen resistance in C. elegans. PLoS Pathog 15: e1007528.

Reid, X. J., J. K. K. Low and J. P. Mackay, 2022 A NuRD for all seasons. Trends in Biochemical Sciences 48: 11–25.

Reynolds, N., P. Latos, A. Hynes-Allen, R. Loos, D. Leaford et al., 2012 NuRD suppresses pluripotency gene expression to promote transcriptional heterogeneity and lineage commitment. Cell Stem Cell 10: 583–594.

Rivera, D. E., V. Lazetic, E. R. Troemel and R. J. Luallen, 2022 RNA Fluorescence in situ Hybridization (FISH) to Visualize Microbial Colonization and Infection in Caenorhabditis elegans Intestines. J Vis Exp 185: e63980.

Rodriguez-Crespo, D., M. Nanchen, S. Rajopadhye and C. Wicky, 2022 The zinc-finger transcription factor LSL-1 is a major regulator of the germline transcriptional program in C. elegans. Genetics 221 :iyac039.

Saudenova, M., and C. Wicky, 2018 The Chromatin Remodeler LET-418/Mi2 is Required Cell Non-Autonomously for the Post-Embryonic Development of Caenorhabditis elegans. J Dev Biol 7: 10.3390/jdb7010001.

Sherman, B. T., M. Hao, J. Qiu, X. Jiao, M. W. Baseler et al., 2022 DAVID: a web server for functional enrichment analysis and functional annotation of gene lists (2021 update). Nucleic Acids Res 50: W216–W221.

Shin, T. H., and C. C. Mello, 2003 Chromatin regulation during C. elegans germline development. Curr Opin Genet Dev 13: 455–462.

Signolet, J., and B. Hendrich, 2015 The function of chromatin modifiers in lineage commitment and cell fate specification. FEBS J 282: 1692–1702.

Solari, F., and J. Ahringer, 2000 NURD-complex genes antagonise Ras-induced vulval development in Caenorhabditis elegans. Curr Biol 10: 223–226.

Sreenivasan, K., A. Rodriguez-delaRosa, J. Kim, D. Mesquita, J. Segales et al., 2021 CHD4 ensures stem cell lineage fidelity during skeletal muscle regeneration. Stem Cell Reports 16: 2089–2098.

Tecle, E., and E. R. Troemel, 2022 Insights from C. elegans into Microsporidia Biology and Host-Pathogen Relationships. Exp Suppl 114: 115–136.

Tecle, E., P. Warushavithana, S. Li, M. J. Blanchard, C. B. Chhan et al., 2025 Conserved chromatin regulators control the transcriptional immune response to intracellular pathogens in Caenorhabditis elegans. PLoS Genet 21: e1011444.

Timmons, L., D. L. Court and A. Fire, 2001 Ingestion of bacterially expressed dsRNAs can produce specific and potent genetic interference in Caenorhabditis elegans. Gene 263: 103–112.

Troemel, E. R., M. A. Felix, N. K. Whiteman, A. Barriere and F. M. Ausubel, 2008 Microsporidia are natural intracellular parasites of the nematode Caenorhabditis elegans. PLoS Biol 6: 2736–2752.

Turcotte, C. A., S. A. Sloat, J. A. Rigothi, E. Rosenkranse, A. L. Northrup et al., 2018 Maintenance of Genome Integrity by Mi2 Homologs CHD-3 and LET-418 in Caenorhabditis elegans. Genetics 208: 991–1007.

Unhavaithaya, Y., T. H. Shin, N. Miliaras, J. Lee, T. Oyama et al., 2002 MEP-1 and a homolog of the NURD complex component Mi-2 act together to maintain germline-soma distinctions in C. elegans. Cell 111: 991–1002.

von Zelewsky, T., F. Palladino, K. Brunschwig, H. Tobler, A. Hajnal et al., 2000 The C. elegans Mi-2 chromatin-remodelling proteins function in vulval cell fate determination. Development 127: 5277–5284.

Wilczewski, C. M., A. J. Hepperla, T. Shimbo, L. Wasson, Z. L. Robbe et al., 2018 CHD4 and the NuRD complex directly control cardiac sarcomere formation. Proc Natl Acad Sci U S A 115: 6727–6732.

Xue, Y., J. Wong, G. T. Moreno, M. K. Young, J. Cote et al., 1998 NURD, a novel complex with both ATP-dependent chromatin-remodeling and histone deacetylase activities. Mol Cell 2: 851–861.

Yoshida, T., I. Hazan, J. Zhang, S. Y. Ng, T. Naito et al., 2008 The role of the chromatin remodeler Mi-2beta in hematopoietic stem cell self-renewal and multilineage differentiation. Genes Dev 22: 1174–1189.

Zaghet, N., K. Madsen, F. Rossi, D. F. Perez, P. G. Amendola et al., 2021 Coordinated maintenance of H3K36/K27 methylation by histone demethylases preserves germ cell identity and immortality. Cell Rep 37: 110050.

Zhang, T., G. Wei, C. J. Millard, R. Fischer, R. Konietzny et al., 2018 A variant NuRD complex containing PWWP2A/B excludes MBD2/3 to regulate transcription at active genes. Nat Commun 9: 3798.

Zhu, D., X. Wu, J. Zhou, X. Li, X. Huang et al., 2020 NuRD mediates mitochondrial stress– induced longevity via chromatin remodeling in response to acetyl-CoA level. Sci Adv: 6, eabb2529.

