## Supplementary figures for "The chromatin remodeler LET-418/Mi-2 regulates the intracellular pathogen response in the *C. elegans* intestine"

**A**

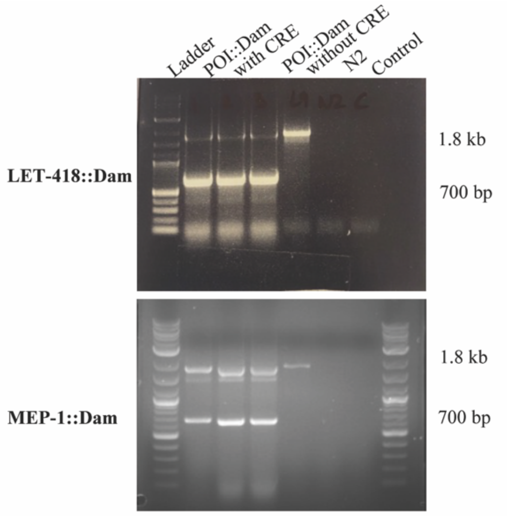


**B**

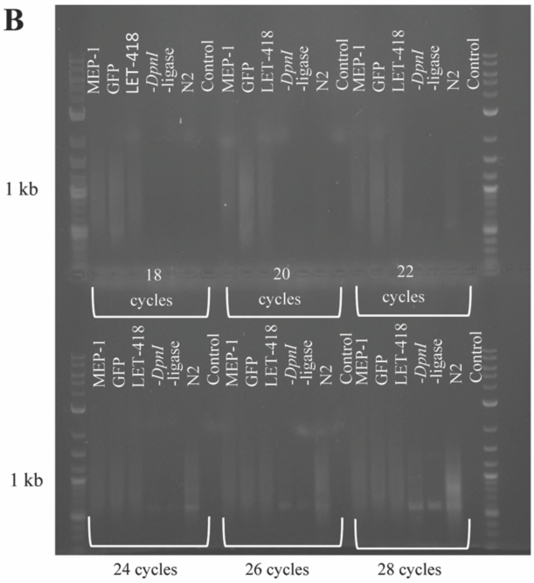


**C**

| Sample | Number of reads |
|---|---|
| MEP1_Intestine_Ya_Rep1 | 1’707’600 |
| MEP1_Intestine_Ya_Rep2 | 2’144’821 |
| LET418_Intestine_Ya_Rep1 | 2’805’532 |
| LET418_Intestine_Ya_Rep2 | 2’355’610 |
| GFP_Intestine_Ya_Rep1 | 842’776 |
| GFP_Intestine_Ya_Rep2 | 775’579 |

**D**

Total number of peaks

| | LET-418 | MEP-1 |
|---|---|---|
| **Rep1** | 5261 | 5635 |
| **Rep2** | 5135 | 5355 |

**E**

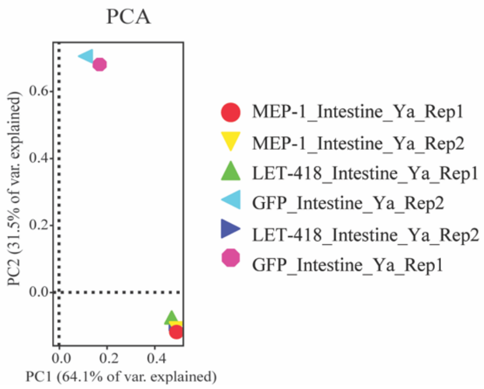


**F**

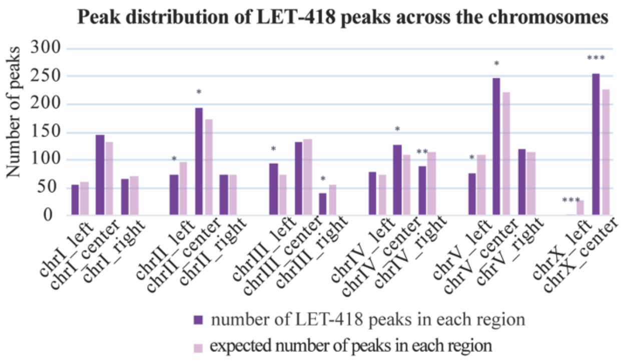


**G**

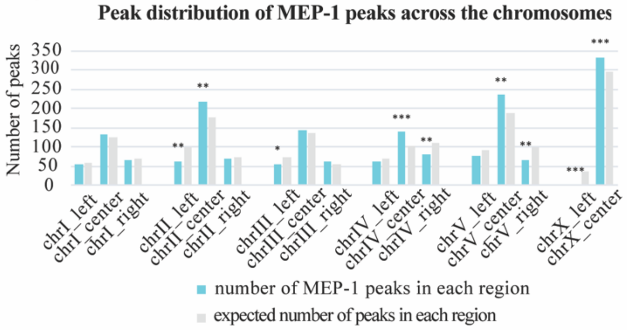


**Sup Figure 1: Specific sequencing of the LET-418 and MEP-1 Dam methylated sequences**
A) To check whether the mCherry cassette is indeed excised from the strains expressing the POI::Dam fusion construct (Protein Of Interest) in the intestine, PCR was performed with primers designed outside the *mCherry* cassette. Since the CRE recombinase is expressed specifically in the intestine, we expect a fragment including the *mCherry* cassette (at 1821 bp) as well as one without the cassette (at 670 bp). Strains without the *CRE* recombinase gene and N2 wild type worms served as negative PCR controls. B) Following *DpnI* digestion of the methylated genome and ligation of adaptors, PCR of the resulting fragments was performed to determine the optimal number of amplification cycles before library preparation and

sequencing. Results are displayed for LET-418 replicate 1. To ensure specific amplification of methylated fragments, control conditions are included. Brief description of the controls: 1) -*DpnI* control includes no *DpnI* enzyme hence no restriction fragments formed. 2) –lig control has no ligase hence no ligation of adaptors. 3) N2 wild type worms do not express the Dam fusion construct. 4) Control is a reaction without any DNA. Samples were analysed after every two cycles from 18 cycles onwards. The optimal amplification resides between 18 to 22 cycles. C) Total number of reads for each replicate resulting from the nanopore sequencing. D) Total number of peaks for each replicate E) Principal Component Analysis (PCA) of the biological replicates resulting from the sequencing of fragments methylated by LET-418::DAM, MEP-1::DAM and GFP::DAM fusion proteins. F-G) Distribution of the LET-418 and MEP-1 binding peaks across the arms and center of chromosomes compared to the expected number of peaks in those regions. Hypergeometric distribution test was performed on the expected and observed number of peaks to know the significance (p-value) between the two values.

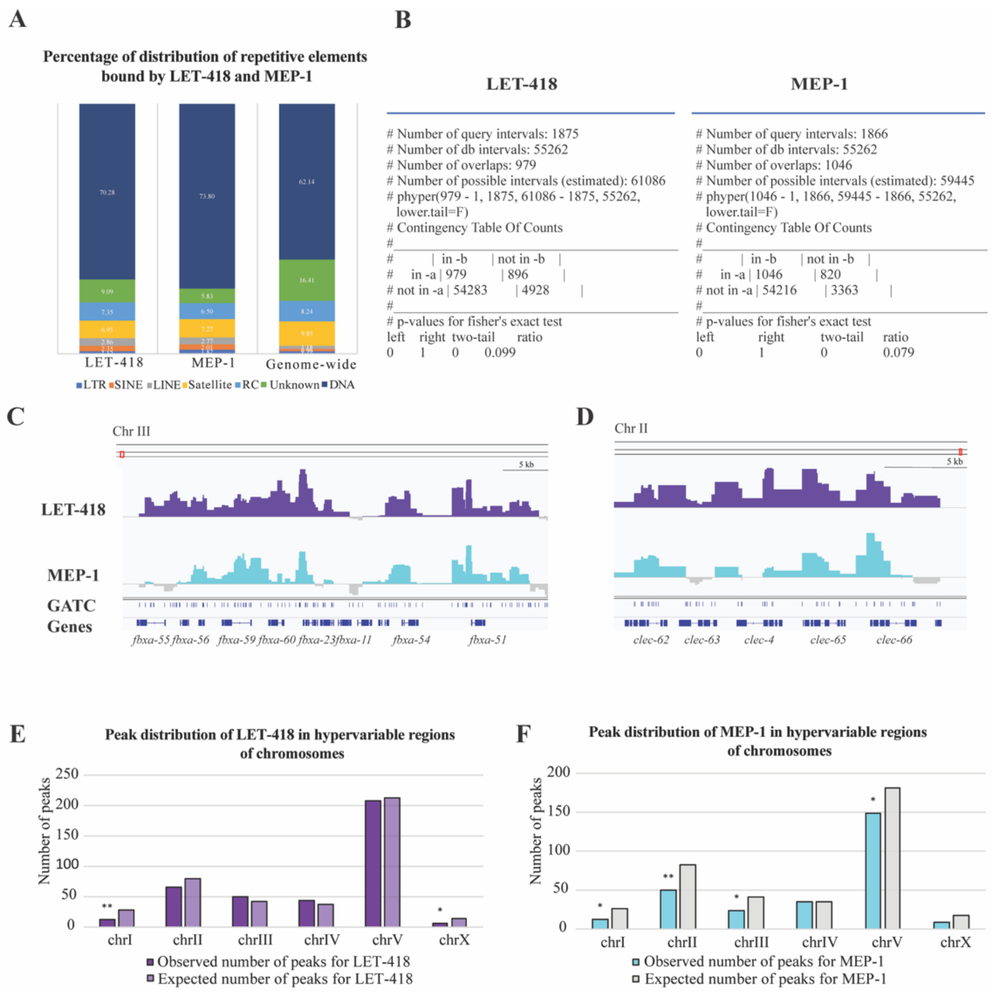


**Sup Figure 2**: **LET-418 and MEP-1 are not binding preferentially to repetitive elements nor to hypervariable regions** A) Repetitive element proportion present in the LET-418 and MEP-1 datasets using stacked plot along with genome wide total distribution of the different repetitive elements. The reference bed file was downloaded from the UCSC browser and edited to exclude simple repeats and low variablity DNA. The resulting file was used to find the proportion of repetitive elements present in the two data sets by using the bedtools intersect function. The output file was analysed using excel to generate the stacked plots. B) Fisher exact test was used to calculate the P value, showing that no significant enrichment in repetitive elements was observed in the LET-418 and MEP-1 binding sites. C) LET-418 and MEP-1 binding to *fbxa* and *clec* gene clusters on Chromosome III and II respectively with a scale of (-0.541-1.71). The *clec* clusters are present in the hypervariable region of chromosome II (LEE *et al.* 2021). Present vs expected peaks in the hypervariable regions for E) the LET-418 and F) the MEP-1 datasets (FDR<0.05). Hypergeometric distribution test was performed on the expected and observed number of peaks to know the significance (p-value) between the two values.

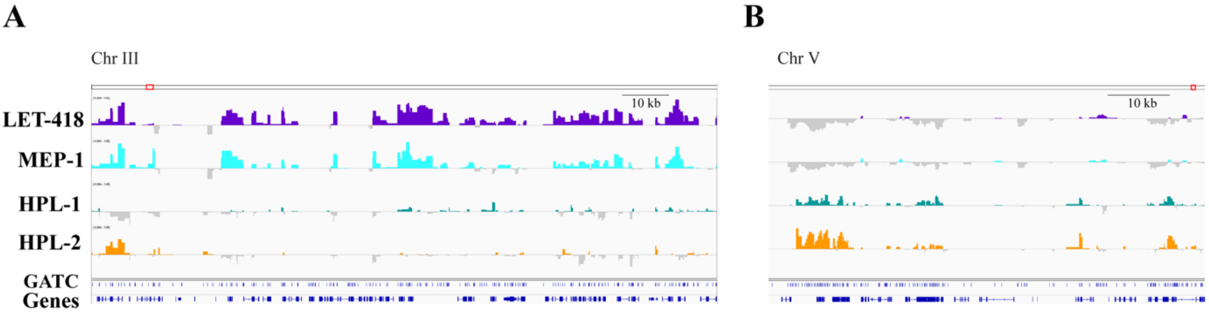


**Sup Figure 3:** IGV browser representations of A) LET-418 and MEP-1 bound regions depleted from HPL-1/2 on chromosome III, scale (-0,954-1,95) and B) HPL-1 and HPL-2 bound regions depleted from LET-418 and MEP-1 on chromosome V, scale (-2,035-2,21) respectively.

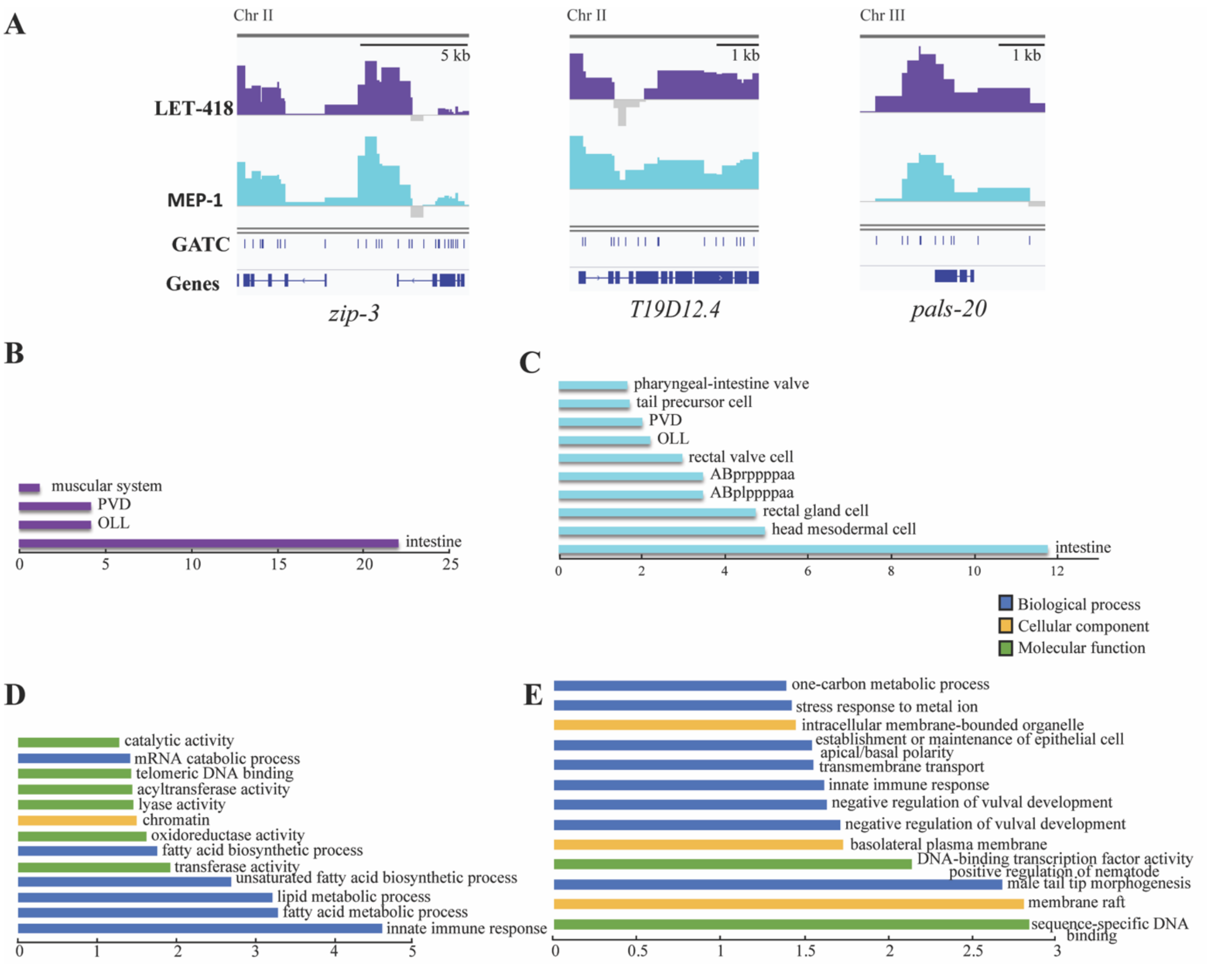


**Sup Figure 4: Target genes unique to either LET-418 or MEP-1 are associated to intestinal functions and innate imuune response** A) Representation on IGV browser of the LET-418 and MEP-1 binding to *zip-3* and *T19D12.4* on Chromosome II, scale of (-0.354-2.08) and (-0.754-1.45) respectively, and *pals-20* on Chromosome III, with a scale of (-0.365-1.72). B,C) Tissue enrichment analysis and D,E) Gene ontology analysis of target genes unique to B,D) LET-418 and C,E) MEP-1 using an intestine specific gene list as a background list (KALETSKY *et al.* 2018). X axis is representing the –log10(p-value).

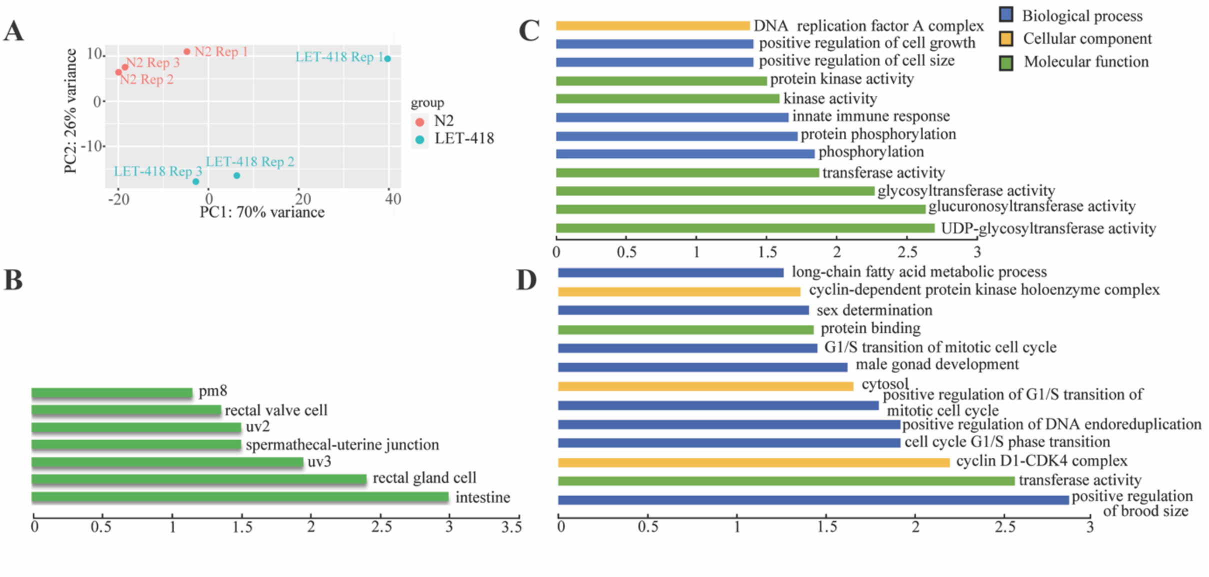


**Sup Figure 5:** LET-418 transcriptome analysis A) Principle Component Analysis of the LET-418 transcriptome data including three replicates for each genotype, i.e. wild type and *let-418* mutant. B) Tissue enrichment analysis of genes that are bound by LET-418 and deregulated in *let-418* mutants. q value threshold of 0.1. X axis is representing the –log10(q-value). Gene ontology using DAVID database of the genes bound by LET-418 and C) up- D) down-regulated in *let-418* mutant. X axis is representing the –log10(p-value).


Kaletsky, R., V. Yao, A. Williams, A. M. Runnels, A. Tadych *et al.*, 2018 Transcriptome analysis of adult Caenorhabditis elegans cells reveals tissue-specific gene and isoform expression. PLoS Genet 14**:** e1007559.

Lee, D., S. Zdraljevic, L. Stevens, Y. Wang, R. E. Tanny *et al.*, 2021 Balancing selection maintains hyper-divergent haplotypes in Caenorhabditis elegans. Nat Ecol Evol 5**:** 794-807.
